# New Tools for Genetically Targeting Myeloid Populations in the Central Nervous System

**DOI:** 10.1101/2019.12.20.885244

**Authors:** Gabriel L. McKinsey, Carlos O. Lizama, Amber E. Keown-Lang, Abraham Niu, Elin Chee, Nicolas Santander, Thomas D. Arnold

**Affiliations:** Department of Pediatrics, University of California San Francisco; Cardiovascular Research Institute, University of California San Francisco; University of California, Berkeley

## Abstract

As the resident macrophages of the brain and spinal cord, microglia are crucial for the phagocytosis of infectious agents, apoptotic cells and synapses. Developmentally, microglia originate from the embryonic yolk sac and serve important roles in the sculpting of neonatal neural circuits. During brain injury or infection, bone-marrow derived macrophages invade neural tissue, making it difficult to distinguish between invading macrophages and resident microglia. In addition to circulation-derived monocytes, other non-microglial central nervous system (CNS) macrophage subtypes include borderzone (meningeal and perivascular) and choroid plexus macrophages. To distinguish between resident microglia and these other CNS macrophage subtypes, we generated a *P2ry12-CreER* mouse line. P2RY12 is a microglial-specific nucleotide sensing GPCR that is important for microglial response to tissue damage. Using immunofluorescent labeling and flow cytometry experiments, we show that *P2ry12-CreER* recombination is exceptionally specific to parenchymal microglia. We also perform ribosome immunoprecipitations and transcriptional profiling of *P2ry12-CreER* recombined cells, using a Cre-dependent *Rpl22-HA* mouse line. By identifying genes enriched in this dataset that are not correspondingly enriched in a broader *Cx3CR1-CreER; Rpl22* dataset, we isolate a number of borderzone macrophage-specific transcripts, including the gene *PF4*. Using a *PF4-Cre* mouse line, we show that *PF4* expression robustly marks borderzone macrophages. Together, we demonstrate two new methods to genetically target distinct CNS macrophage subtypes.

## Introduction

CNS macrophages can be broadly separated into three types: microglia, choroid plexus macrophages, and borderzone macrophages of the dura, subdura and the perivascular space. Circulatory monocytes, which invade the CNS and differentiate into macrophages, are also present in the context of CNS injury and inflammation. These different types of macrophages play fundamental roles in CNS development, homeostasis and disease, but their specific functions remain poorly defined. One primary barrier to defining the subtype-specific functions of CNS macrophages has been a lack of genetic tools to separately target these subpopulations.

Microglia, the resident macrophages of the neural parenchyma, regulate a wide variety of processes in the brain, from development and synapse remodeling, to inflammatory insult and antigen presentation. Fate-mapping and developmental analyses have revealed that microglia are derived from the embryonic yolk sac, unlike circulating monocytes, which are derived from the adult bone marrow. Although bone-marrow derived macrophages can adopt some features of endogenous microglia, they are not able to fully recapitulate all of their properties, suggesting that there may be important functional differences between these two populations of cells. The use of single-cell transcriptomic profiling has revealed new insights about microglial biology in recent years (Hammond et al., 2019; Jordão et al., 2019; Li et al., 2019; Van Hove et al., 2019) suggesting a greater degree of heterogeneity among microglia than previously recognized. Recently developed microglial genetic labeling strategies, such as the mouse line *Cx3cr1-CreER* (Goldmann et al., 2013a; Parkhurst et al., 2013; Yona et al., 2013), have significant advantages compared to their predecessors, but still have a number of drawbacks. For instance, *Cx3cr1* is expressed in other immune cell types in the brain, including perivascular, choroid plexus and meningeal macrophages (Goldmann et al., 2016). *Cx3cr1-CreER* also labels circulating monocytes, which results in the labeling of peripherally-derived cells in the context of CNS injury or disease. Furthermore, *Cx3cr1-CreER* mouse lines are haplo-insufficient for *Cx3cr1*, which may affect microglia functions, such as synaptic pruning (Rogers et al., 2011).

Here, we describe a *P2ry12-CreER* knock-in mouse line that we have developed to specifically label brain microglia. P2RY12 is a nucleotide sensing metabotropic GPCR in the P2Y family of GPCRs that has an important role in the microglial “sensome” (Hickman et al., 2013). P2RY12 regulates the microglial response to tissue damage and the adoption of the amoeboid “activated” microglial morphology (Bernier et al., 2019; Haynes et al., 2006). We found that *P2ry12-CreER*, unlike the commonly used *Cx3cr1-CreER*, specifically labeled brain microglia, but not perivascular or meningeal macrophages. Using this line, we performed ribosomal immunoprecipitations and transcriptional profiling of “pure” microglia. By comparing this dataset to existing ribosomal profiles of CNS macrophages and subtracting genes that are shared by both datasets, we identify a number of borderzone macrophage markers, including *PF4*. Analysis of the *PF4-Cre* mouse line shows that *PF4*-*Cre* recombination robustly labels perivascular and meningeal macrophages. Altogether, we show that *P2ry12-CreER* and *PF4-Cre* can be used to genetically target parenchymal microglia and perivascular macrophages, two distinct CNS macrophage subtypes.

## Results

### Generation of a P2ry12-CreER mouse line

To identify specific markers of brain microglia, we cross-referenced published reports of microglial markers (Gautier et al., 2012) (Buttgereit et al., 2016) (Wieghofer et al., 2015) with recently produced single cell sequencing data sets (Tabula Muris: https://tabula-muris.ds.czbiohub.org, Myeloid Cell Single Cell Seq database: https://myeloidsc.appspot.com). We analyzed several microglial markers (*HexB, P2ry12, Sall1, Tmem119, Trem2, Fcrls* and others), and compared their expression across cell types in the mouse body. Among these genes, *P2ry12* appeared to be the most restricted to brain myeloid cells.

To genetically label P2RY12(+) microglia, we generated a *P2ry12-2A-CreER* knock-in mouse line (hereafter called *P2ry12-CreER*) **(Figure 1A)** using CRISPR-facilitated homologous recombination. The targeting construct was designed to preserve *P2ry12* expression, with the *P2ry12* stop codon replaced by a ribosome-skipping 2A-fusion peptide coding sequence, followed by the coding sequence for *CreER*. We did not find any health problems in either *P2ry12-CreER* heterozygous or homozygous mice. Immunolabeling of homozygous and heterozygous *P2ry12-CreER* mice showed no significant change in P2RY12 protein expression or microglial morphology in homozygous compared to wild-type mice **(Fig 1A)** indicating that gene knock-in did not affect gene expression or function.

**Figure 1.**
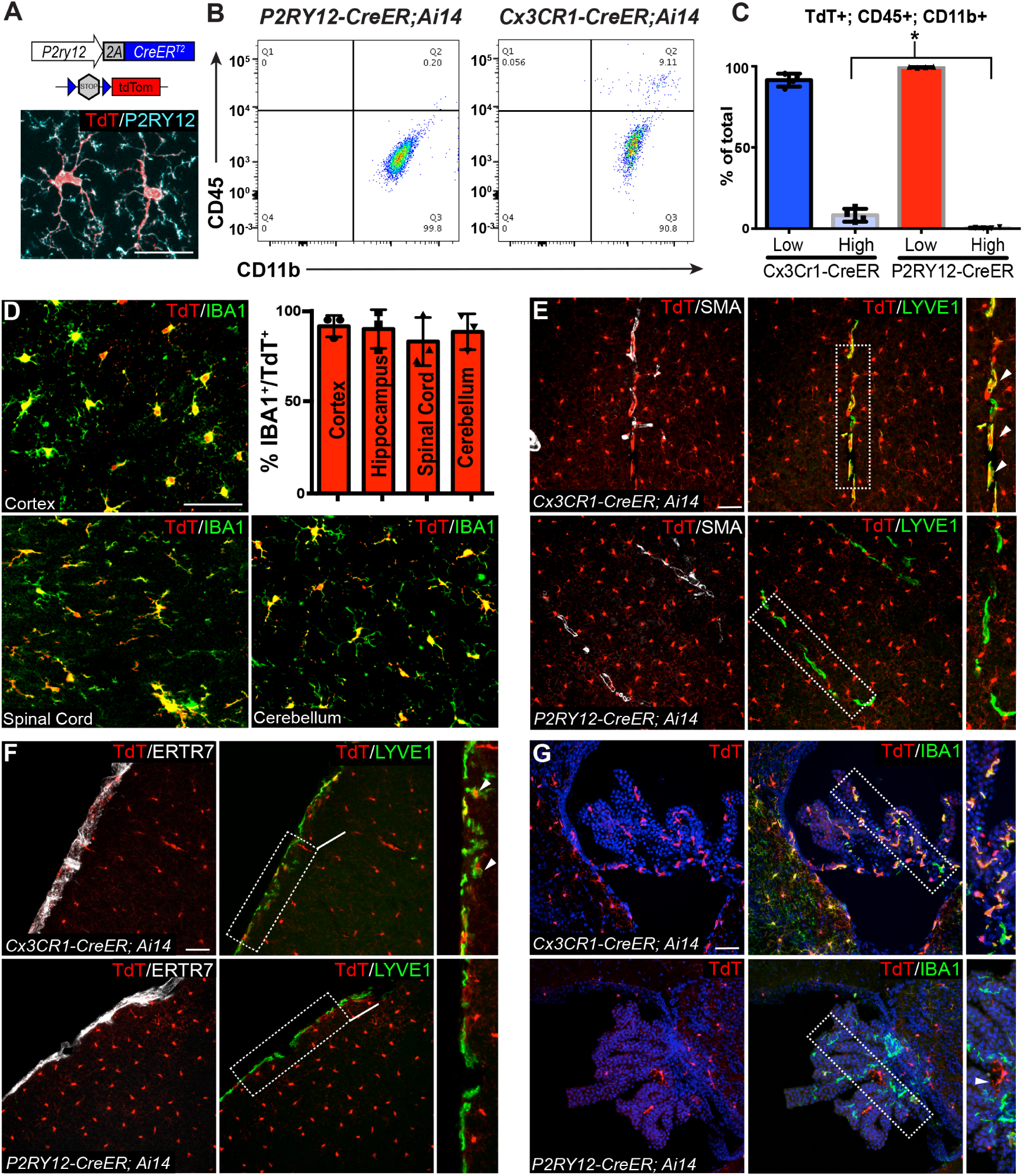
Microglia are specifically and efficiently recombined by *P2ry12-CreER*. **A**. Design of the *P2ry12* knock-in allele. Brain section from *P2ry12-CreER;Ai14* reporter mouse stained for microglia (P2RY12, cyan) and recombined cells (TdTomato, red). **B**. Flow cytometry analysis of recombined TdTomato+ macrophages in *P2ry12-CreER;Ai14* and *Cx3cr1-CreER;Ai14* mice. TdTomato+ cells were pre-gated on forward scatter, isolation of singlets, and removal of dead cells. **C**. Quantification of flow cytometry analysis. **D**. Images and quantification of recombination in the cerebral cortex, hippocampus, cerebellum and spinal cord in *P2ry12-CreER;Ai14* mice. Sections stained with pan-macrophage marker IBA1 (green). **E**. Analysis of *Cx3cr1-CreER* vs *P2ry12-CreER* mediated recombination in aSMA-adjacent, LYVE1(+) perivascular macrophages. *Cx3cr1-CreER* recombination labels perivascular macrophages (arrowheads), but *P2ry12-CreER* does not. **F**. Analysis of *Cx3cr1-CreER* vs *P2ry12-CreER* mediated recombination in LYVE1(+) meningeal macrophages. *Cx3cr1-CreER* recombination labels meningeal macrophages (arrowheads), but *P2ry12-CreER* does not. ERTR7 expression delineates meningeal boundaries. **G**. Analysis of *Cx3cr1-CreER* vs. *P2ry12-CreER* mediated recombination in the choroid plexus. *Cx3cr1-CreER* recombination labels most IBA1(+) choroid plexus macrophages; *P2ry12-CreER* recombination labels presumptive Kolmer epiplexus cells on the surface of the choroid plexus (arrowhead). Sections stained with pan-macrophage marker IBA1 (green), vascular marker aSMA (white), meningeal fibroblast marker (ERTR7, white) and meningeal macrophage marker LYVE1 (green). N = 4 separate animals for each genotype in C. and 3 animals for analysis in D. For flow analysis in C., p=.032, Student’s t-test. Scale Bar = 50um (D,E,F,G).

### P2ry12-CreER-mediated recombination efficiently and specifically labels microglia, but not CNS border-associated macrophages

To determine the efficiency and specificity of *P2ry12-CreER* recombination, we generated *P2ry12-CreER*; *Ai14* reporter mice, which express TdTomato upon Cre-dependent recombination(“A robust and high-throughput Cre reporting and characterization system for the whole mouse brain,” 2009). Following tamoxifen induction in adult mice, we saw robust TdTomato expression in P2RY12^+^ microglia **(Fig 1A)**. Analysis of *P2ry12-CreER* recombination in the brain by flow cytometry showed that CD11b^+^CD45^low^ microglia but not CD11b^+^CD45^high^ non-microglial macrophages were preferentially marked by *P2ry12-CreER* recombination. 99.55% and .42% of TdTomato^+^ cells were CD11b^+^; CD45^low^ and CD11b^+^; CD45^high^, respectively, in *P2ry12-CreER; Ai14* mice. In comparison, 91.45% and 8.26% of TdTomato^+^ cells were CD11b^+^; CD45^low^ and CD11b^+^; CD45^high^, respectively, in *Cx3cr1-CreER; Ai14* mice **(Figure 1B,C)**, suggesting that *P2ry12-CreER* recombination is much more specific for parenchymal microglia than *Cx3cr1-CreER*, which recombines many CNS macrophage subpopulations, including perivascular, meningeal and choroid plexus macrophages. A systematic immunohistochemical analysis of *P2ry12-CreER; Ai14* recombination showed high levels of microglial recombination throughout the CNS, as measured by overlap with the pan-macrophage marker IBA1 **(Figure 1D)**.

Consistent with our flow cytometry results, we saw no recombination in perivascular macrophages (LYVE1^+^ macrophages associated with αSMA immuno-labeled vessels), unlike *Cx3cr1-CreER*, which robustly labels these cells **(Figure 1E)**. Similarly, we saw no recombination in the meninges of *P2ry12-CreER; Ai14* mice, unlike *Cx3cr1-CreER; Ai14* mice, which showed recombination in a significant portion of LYVE1^+^ meningeal macrophages. In the choroid plexus, *P2ry12-CreER* recombination marked a small number of IBA1^+^ cells, as well as a small number of IBA1-negative or IBA1-low cells, unlike *Cx3cr1-CreER*, which showed recombination in most choroid plexus IBA1^+^ macrophages **(Figure 1G)**. Some of the *P2ry12-CreER; Ai14* recombined cells appeared to be on the surface of the choroid plexus, consistent with a recently published study that revealed a microglia-like, *P2ry12*-expressing “CP-epi” subpopulation of choroid plexus macrophages, which may be Kolmer’s epiplexus cells (Ling et al., 1998; Van Hove et al., 2019).

### P2ry12-CreER recombination efficiently and specifically labels embryonic microglia

In our previous studies, we noted that P2RY12 expression is developmentally regulated, increasing in the brain throughout embryonic and early post-natal development (Arnold et al., 2019). To determine whether *P2ry12-CreER* was capable of recombining embryonic macrophages, we induced pregnant dams carrying *P2ry12-CreER; Ai14* pups with a series of three tamoxifen doses at E13.5, E15.5, and E17.5 **(Figure 2A)**. We collected these pups at E18.5 and analyzed their degree and pattern of recombination. Examination of these brains revealed widespread recombination in IBA1^+^ parenchymal microglia. Recombination was seen in essentially all parenchymal IBA1^+^ cells, which were most dense in the subventricular zone and lateral migratory stream of the amygdala, where at this age there are significant numbers of migrating GABAergic interneurons **(Figure 2B-B’’, C, D)**. Unlike the widespread parenchymal IBA1^+^ recombination, we observed sparse recombination in the embryonic meninges that was limited to LYVE1^+^ macrophages **(Figure 2C, E)**. Embryonic recombination in IBA1^+^ cells of the choroid plexus was widespread, unlike in the adult **(Figure 2F)**. We did not observe any embryonic recombination in blood vessels or other cell types in the brain at this age.

**Figure 2.**
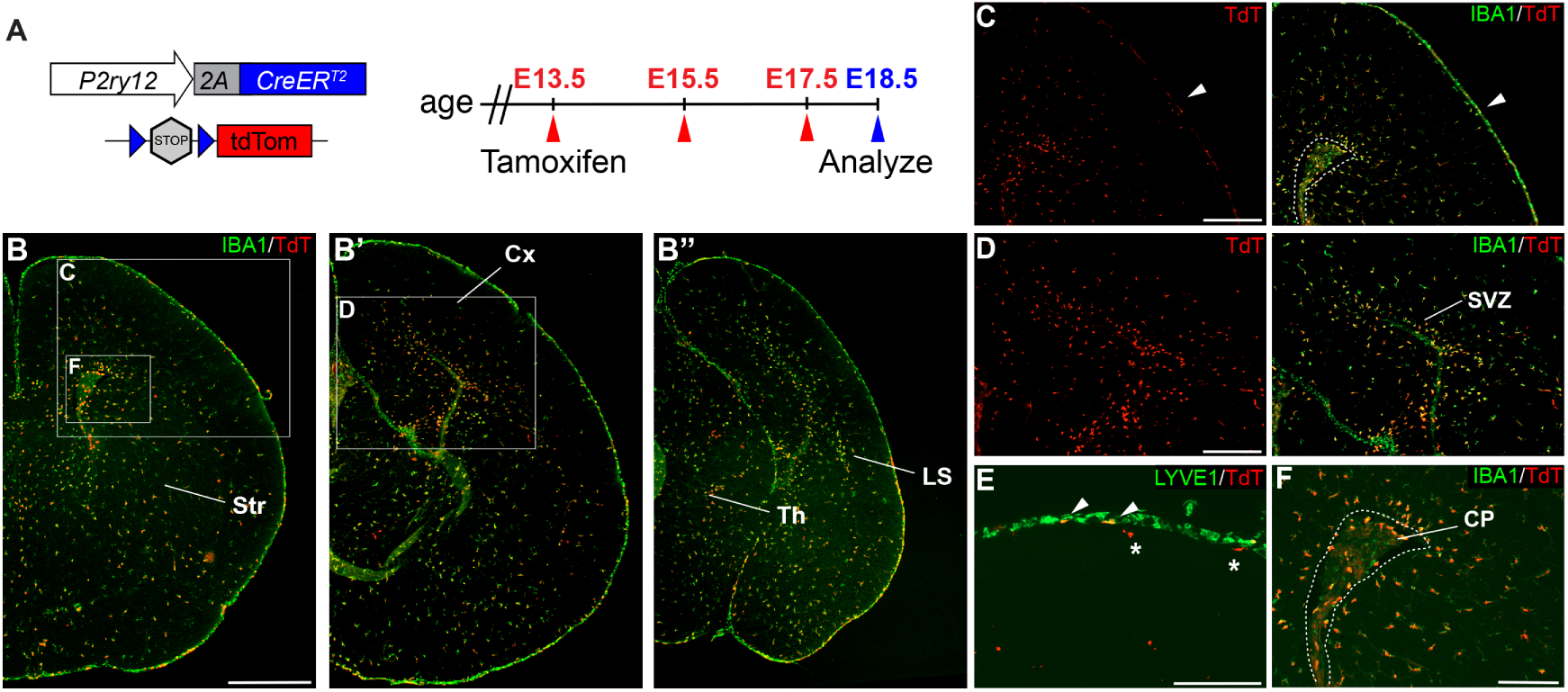
Embryonic recombination in P2ry12-CreER mice. **A**. Diagram of *P2ry12-CreER; Ai14* embryonic tamoxifen induction regimen. Pregnant mice were induced with tamoxifen at E13.5, E15.5 and E17.5 and embryos were collected at E18.5. **B**. Recombination in the E18.5 brain, shown in three images, arranged rostral (B) to caudal (B’’). **C**. *P2ry12-CreER; Ai14* recombination in the developing cerebral cortex. Recombination was seen primarily in the parenchyma of the brain, with sparse recombination in the meninges (arrowhead). **D**. Recombination in the developing hippocampus. **E**. Sparse recombination of LYVE1-expressing meningeal macrophages. **F**. Recombination in the choroid plexus was more widespread at E18.5, marking the majority of IBA1(+) macrophages. Cx=Cortex, LS=Lateral migratory stream, Str=Striatum, Th=Thalamus, SVZ=Subventricular zone. Scale bars= 800um (B), 400um (C, D), 200um (E, F).

### Expression of P2ry12-CreER in other tissues and in circulating monocytes

To determine the specificity of *P2ry12-CreER*-mediated recombination, we examined recombination in non-microglial cell types in the brain, circulation and in a variety of organs. In the brain, we saw no recombination in macroglia (oligodendrocytes and astrocytes), or neurons **(Figure 3A)**. In organs of *P2ry12-CreER*; *Ai14; Cx3Cr1-eGFP* mice, we found sparse recombination that often overlapped with EGFP expression, suggesting *P2ry12* expression mainly marks a small subset of macrophages in other tissues **(Figure 3B)**. Sparse recombination was seen in all organs analyzed (spleen, intestine, heart, skin, lung, liver, muscle, thymus, spleen), except for the kidney, which did not show any recombination (not shown). Strong splenic recombination was seen in a subset of *Cx3cr1-eGFP*^*+*^ cells of the marginal zone between the red and white pulp, an area that contains macrophages that surveil the bloodstream (Borges da Silva et al., 2015). In the lung, intestine and heart, *P2ry12-CreER; Ai14* recombination occurred in a subset of *Cx3cr1-eGFP* cells; in the liver and thymus, *P2ry12-CreER; Ai14* recombination partially overlapped with *Cx3cr1-eGFP*.

**Figure 3.**
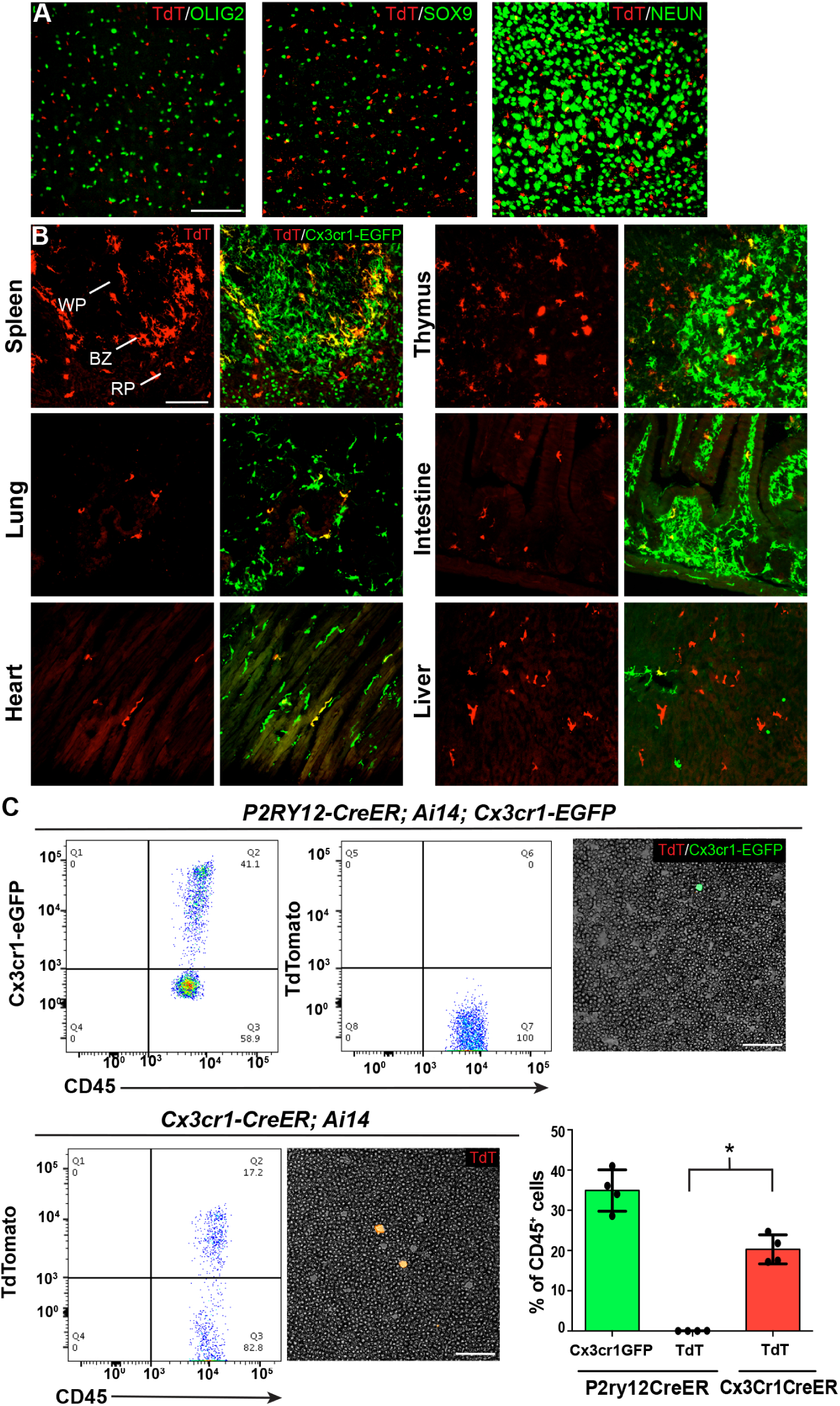
P2ry12-CreER recombination in non-microglial populations is limited. **A**. Analysis of *P2ry12-CreER* recombination showed a lack of recombination in oligodendrocytes (OLIG2), astrocytes (SOX9) or neurons (NEUN). **B**. Analysis of recombination in the spleen, lung, heart, thymus, intestine and liver of *P2ry12-CreER; Ai14; Cx3cr1-eGFP* mice. *Cx3cr1-eGFP* was used as a marker of tissue resident macrophages. **C**. Flow cytometry and blood smear recombination analysis reveals negligible recombination of blood cells in *P2ry12-CreER;Ai14* mice compared to *Cx3cr1-eGFP* (internal control) and *Cx3cr1-CreER;Ai14* mice. p=.0015, Student’s t-test. N = 4 mice for A. and C., 3 mice for B. Scale bars: 100um (A,B,C). WP=White pulp, BZ=Borderzone, RP=Red pulp.

To determine whether *P2ry12-CreER* recombines circulating monocytes, we examined recombination in the circulating blood in smears and by flow cytometry. Blood smears of tamoxifen induced *P2ry12-CreER; Ai14; Cx3Cr1-eGFP* mice showed no recombination in circulating lymphocytes, but as an internal positive control, these mice did have numerous Cx3cr1-EGFP^+^CD45^+^CD11b^+^ circulating monocytes **(Figure 3C)**. In contrast, smears of *Cx3cr1-CreER; Ai14* mice, showed TdTomato expression indicative of recombination in circulating monocytes. Flow cytometry of *P2ry12-CreR; Ai14; Cx3Cr1-eGFP* mice showed a lack of TdTomato+ circulating monocytes (3 samples with 0 TdTomato^+^ cells, 1 sample with 2 TdTomato^+^ cells out of 2852 CD45^+^CD11b^+^ monocytes), whereas *Cx3cr1-CreER; Ai14* samples showed recombination in 20.3% of CD45^+^CD11b^+^ monocytes **(Figure 3C)**. These data indicate that *P2ry12-CreER* recombination labels sparse subsets of resident macrophages in organs other than the brain, but does not label circulating monocytes.

### P2ry12-CreER-dependent ribosomal profiling enriches for transcripts of true brain microglia

We next studied the mRNA transcriptional profile of true microglia using *P2ry12-CreER* mice. Classically, mRNA profiling of defined cell types requires preparation of single cell suspensions followed by cell sorting, RNA isolation and sequencing. These manipulations are inefficient and introduce bias, especially in macrophage populations, which are difficult to mechanically or enzymatically separate. Furthermore, microglia become activated and show significant transcriptional changes upon dissociation and sorting (Haimon et al., 2018a). To avoid these pitfalls, we performed TRAP-Seq (translating ribosome affinity purification followed by mRNA sequencing) (Sanz et al., 2009) which allows for cell-specific isolation of mRNA without these manipulations. We generated *P2ry12-CreER*; *Rpl22-HA* mice, which, upon recombination, express a hemagglutanin (HA) epitope tagged RPL22 riboprotein that can be isolated by immunoprecipitation **(Figure 4A)**. Upon tamoxifen-induced recombination in *P2ry12-CreER; Rpl22-HA* mice, we found robust labeling of microglia by Rpl22-HA, as shown by immunofluorescent costaining for HA and IBA1 **(Figure 4B)**. qPCR analysis of immunoprecipitations from *P2ry12-CreER; Rpl22-HA* mice revealed significant enrichment for microglial transcripts such as *Tmem119, P2ry12* and *HexB (not shown)*. After this qPCR validation, libraries from *P2ry12-CreER; Rpl22* immunoprecipitations were generated and sequenced **(workflow in 4C)**. Analysis of the transcripts enriched in *P2ry12-CreER; Rpl22* libraries revealed significant enrichment for many known microglial markers, but did not show enrichment for markers of other cell types, such as neurons, oligodendrocytes and astrocytes **(Figure 4D)**.

**Fig. 4.**
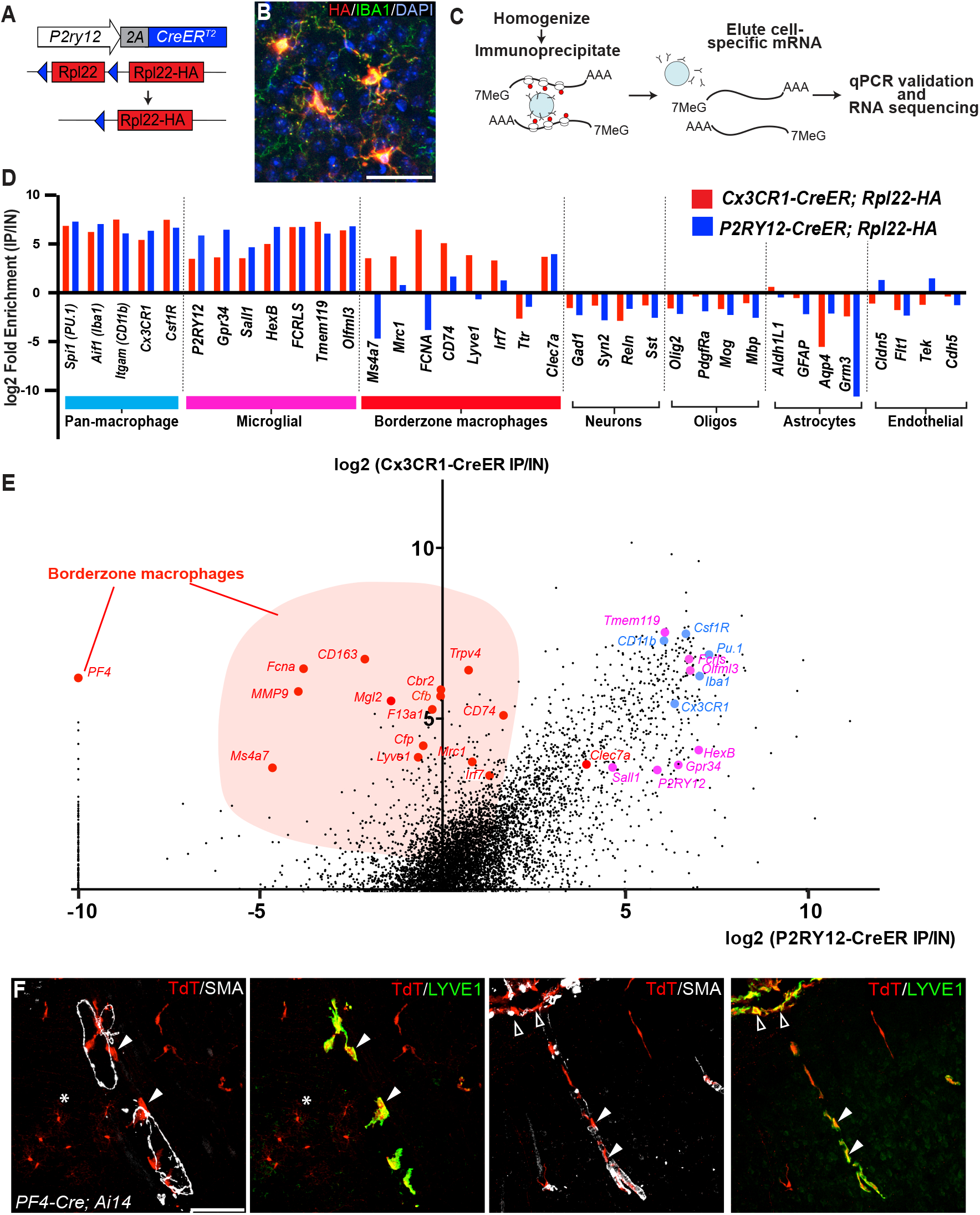
Cre-dependent ribosomal profiling of microglia. **A**. A diagram illustrating *P2ry12-CreER*-dependent recombination of the *Rpl22-HA* allele, used to perform cell-type specific ribosomal profiling. **B**. Immunostaining of tagged microglia for HA (red) and IBA1 (green) shows strong Cre-dependent expression of *Rpl22-HA* in microglia. **C**. Workflow diagram for Cre-dependent ribosomal profiling experiments. **D**. A comparison of relative enrichment for cell-type specific markers from *P2ry12-CreER; Rpl22-HA* and *Cx3cr1-CreER; Rpl22-HA* ribosomal immunoprecipitations. **E**. Genome-wide comparisons of relative enrichment following immunoprecipitation of ribosomes in *P2ry12-CreER; Rpl22-HA* and *Cx3cr1-CreER; Rpl22-HA* mice. Pan-macrophage markers in blue; presumptive microglial markers in pink; presumptive borderzone macrophage markers in red. **F**. *PF4-Cre* robustly recombines perivascular SMA-adjacent LYVE1(+) perivascular macrophages (closed arrowheads) and meningeal macrophages (open arrowheads). Asterisk: recombination in a small cluster of microglia. Scale bars: 40um (B) and 100um (F).

The transcriptional profile of *P2ry12-CreER* recombined cells was quite similar to recently published profiles derived from *Cx3cr1-CreER; Rpl22* immunoprecipitations (Haimon et al., 2018b). By comparing the relative enrichment values for individual genes in our *P2ry12-CreER; Rpl22* data **(X axis, Figure 4E)** to *Cx3cr1-CreER; Rpl22* enrichment values for the same gene **(Y axis, Figure 4E)**, we identified a number of enriched genes that were either common or unique to each dataset. Genes common to both datasets included macrophage markers such as *Pu*.*1, Aif1 (which codes for IBA1), Csf1r* and *CD11b*. In line with our histological and cell-sorting data however, there were many genes enriched in *Cx3cr1-CreER; Rpl22* IPs that were not enriched in *P2ry12-CreER; Rpl22* IPs. Some of these genes have recently been identified in single-cell sequencing studies as markers of border-associated macrophages (Van Hove et al., 2019), suggesting that our comparison differentiates genes expressed in true microglia from those expressed in non-parenchymal macrophages in the CNS. This subtractive approach also identified a number of genes that were preferentially enriched in the *P2ry12-CreER; Rpl22* population, which represent potential microglial specific markers. As expected, *P2ry12* was in this group, as were a number of other *P2Y* and *P2X* family members, such as *Gpr34, P2ry10b, 13, 6* and *P2rx1, 7*.

One gene that had significant enrichment in *Cx3cr1-CreER; Rpl22* IPs but marked depletion in *P2ry12-CreER; Rpl22* IPs was the gene *PF4*, also known as *Cxcl4* **(Figure 4E)**. Given its strong enrichment in *Cx3cr1-CreER* expressing cells and lack of enrichment *P2ry12-CreER* expressing cells, *PF4* appeared to be a good candidate marker of perivascular and/or meningeal macrophages. We explored the lineage of cells marked by *PF4* expression by examining cells recombined by a *PF4-Cre* transgenic mouse line (Tiedt et al., 2007). Indeed, we found that in *PF4-Cre; Ai14* mice, perivascular and meningeal macrophages are strongly recombined **(Figure 4F)**. *PF4-Cre; Ai14* mice also had a low degree of microglial recombination, which appeared as clumps of cells scattered throughout the brain, suggestive of sporadic recombination and subsequent clonal expansion.

## Discussion

In recent years, knock-in and transgenic mouse lines using the fractalkine receptor gene *Cx3cr1* (e.g. *Cx3cr1-Cre*, -*CreER* and -*eGFP*) have been some of the best and most frequently used tools for targeting microglia (Goldmann et al., 2013b; Parkhurst et al., 2013). Other strategies, targeting genes such as *Tie2 (Cre, CreMERM), CD11b (Cre, L10aGFP), and CSF1R (LacZ, eGFP, CreMERM)* (M. L. Bennett and F. C. Bennett, 2019; Wieghofer et al., 2015) have been used in landmark studies to determine the yolk-sac origin of microglia, and to characterize the transcriptional architecture of these cells. However, despite their usefulness, many of these microglia-targeting mouse lines mark other myeloid cell types in the CNS. For instance, *Cx3cr1*-based mouse lines mark borderzone and choroid plexus macrophages in addition to microglia. *Cx3cr1* is also expressed in circulating T cells, B cells, NK cells and monocytes, which means that in the context of injury, *Cx3CR1-Cre* and *eGFP* are incapable of distinguishing between resident macrophages and other invading immune cells (Bar-On et al., 2010; Fogg et al., 2006; Imai et al., 1997; Liu et al., 2009). The tamoxifen-inducible *Cx3cr1-CreER* allele avoids this problem to a degree, but requires a significant waiting period after the time of recombination for recombined circulating cells to be depleted. This waiting period is inconvenient for most models of CNS injury and is incompatible with time-sensitive studies, such as models of neonatal brain injury. In addition to issues regarding specificity, *Cx3cr1* knock-in alleles (*eGFP, Cre, and CreER*) (Goldmann et al., 2013a; Parkhurst et al., 2013) disrupt the *Cx3cr1* gene, which may result in *Cx3cr1*-dependent effects on synaptic pruning (Rogers et al., 2011). Aside from *Cx3CR1*, knock-in *GFP* and *CreER* lines have been generated for the microglial-enriched transcription factor *Sall1* (Buttgereit et al., 2016). However, significant caveats for these lines exist, including the fact that *Sall1-GFP* and *CreER* both disrupt the function of the *Sall1* gene, and that *Sall1-CreER* marks a broad range of non-myeloid CNS cell types (Chappell-Maor et al., 2019).

Here, we show by immunohistochemistry, flow cytometry, and ribosomal profiling that *P2ry12-CreER* has a number of advantages over existing microglial targeting *Cre* and *CreER* mouse lines. One such advantage is that *P2ry12-CreER*-recombination avoids circulating monocytes, meaning that in the context of injury, resident microglia can be distinguished from invading monocyte-derived macrophages. *P2ry12-CreER* also avoids targeting borderzone macrophages of the meninges or perivascular space. As perivascular inflammation is a hallmark of many neurological diseases, such as multiple sclerosis and Alzheimer’s disease, the ability to separately manipulate microglial and borderzone macrophages in mouse models of these diseases will be critical.

The *P2ry12-CreER* mouse line is particularly well-suited for the labeling and tracking of microglia during CNS brain injury. By genetically labeling microglia, transcriptional profiling can be performed at different stages of CNS injury, from the initial inflammatory response to later regenerative stages, without contamination by invading peripheral monocytes. One major barrier to studying microglia in the context of injury is that microglia undergo significant transcriptional changes when activated, making it difficult to distinguish activated microglia from other macrophage subtypes. For instance, upon activation, microglia have been shown to downregulate “true” microglia markers (*Tmem119, P2ry12*) and upregulate other macrophage markers (*CD45, CD86, Cd53, TLR4, TLR7*) (Haimon et al., 2018b; Jordão et al., 2019). Microglial activation occurs in response to CNS injury, but is also induced by methods used to prepare microglia for analysis by flow cytometry. By genetically labeling and profiling microglia with *P2RY12-CreER* and TRAP-seq, these transcriptional changes associated with flow cytometry can be avoided (Haimon et al., 2018a).

The recent publication of a number of *Tmem119*-targeted mouse alleles (*Tmem119-GFP, RFP, CreER*) serve to highlight some of the advantages of the *P2ry12-CreER* line described here (Kaiser and Feng, 2019; Ruan et al., 2020). Regarding the specificity of *Tmem119*-based lines, both *Tmem119-CreER* and –*GFP* appear to show low but not insignificant levels of recombination or expression in brain-derived CD11b+; C45int/hi myeloid cells (1.9% and 6.9% for *CreER* recombination and GFP expression respectively). In the circulating blood, no *Tmem119-GFP*(+) monocytes were reported, but 7 days after the last tamoxifen injection, 3% of circulating CD45(+) monocytes were labeled by *Tmem119-CreER*. In a separate publication, using a Tmem119-RFP mouse line and a percoll gradient to isolate CNS macrophages, 5.3% of RFP+ cells of CNS macrophages were shown to be CD11b^+^; CD45^hi^, similar to the Tmem119-GFP report. In the neonatal P1 brain, *Tmem119-GFP* showed widespread expression overlapping with the vascular endothelial cell marker, CD31. In adulthood, prolonged administration of tamoxifen to *Tmem119-CreER* mice induced recombination in non-IBA1(+) cells. This recombination was quite strong in IBA1(-) meningeal cells and in cells adjacent to large penetrating blood vessels, likely representing meningeal and perivascular fibroblasts, respectively. Indeed, based on single cell sequencing, Tmem119 is highly expressed in CNS resident fibroblasts (http://betsholtzlab.org/VascularSingleCells/database.html). In summary, despite robust labeling of microglia in *Tmem119*-based lines, recombination in CD11b+; C45int/hi CNS cells and circulating monocytes by flow cytometry, the recombination in mural and meningeal cells, and the significant degree of *Tmem119-GFP* endothelial expression in neonatal brains suggest that *P2ry12* may be a more useful gene for genetic targeting of microglia.

*P2ry12-CreER* can be used to delete floxed alleles in embryonic or adult microglia. Here, we show that in addition to the fluorescent TdTomato-expressing Ai14 reporter mouse line (Madisen et al., 2009), *P2ry12-CreER* can be used to recombine the *Rpl22-HA* ribotrap allele for microglia-specific transcriptional profiling. Interestingly, using a subtractive approach, we found a number of genes that had very high enrichment in *Cx3cr1-CreER; Rpl22* samples but no enrichment in *P2ry12-CreER; Rpl22* samples. Some of these markers (*Mrc1, F13a1, Ms4a7* and *MMP9*) were recently identified in other studies that used physical dissociation of the dural membranes from the neural parenchyma to separate borderzone macrophages from other CNS macrophage subtypes (Jordão et al., 2019; Van Hove et al., 2019). After isolating macrophages from these preparations, these groups performed single-cell sequencing to identify gene signatures of microglia, border-associated and choroid plexus macrophages. One major drawback to strategies that rely on the physical dissection of these populations is that this process results in significant levels of cross contamination from microglia and other cells that that are in close contact with the meningeal layers. Furthermore, physical dissection of the meningeal layers does not separate perivascular macrophages of the penetrating vessels from the parenchymal microglia. In contrast, *P2ry12-CreER* recombination provides a more precise and complete genetic segregation of these populations that minimizes the transcriptional changes that are known to occur during sorting. The differential enrichment observed in our subtractive approach revealed several transcripts only expressed in border-associated macrophages. We focused on one such gene, *PF4*, and found that *PF4-Cre* robustly and preferentially recombines borderzone macrophages. We did observe a low degree of sporadic microglial recombination in *PF4-Cre; Ai14* mice, which may be due to short-term expression of *PF4*, perhaps upon microglial activation, or developmental labeling. Together, our studies find that with *P2ry12-CreER and PF4-Cre*, both parenchymal and borderzone CNS macrophages can be genetically dissociated.

CNS macrophages regulate a wide range of processes from early development to adulthood in health and disease. Here, we provide improved methods for genetically targeting microglia and borderzone macrophages. Use of these genetic tools will open new experimental avenues to better define the specific functions of CNS macrophage subclasses. For instance, microglial or borderzone macrophages ablation experiments can be performed with conditional *Csf1R* ablation, or conditional diphtheria toxin receptor expression. Another particularly useful function of these lines will be to track and transcriptionally profile distinct macrophages classes in the context of CNS injury or neurodegenerative disease.

## Material and Methods

### Mice

*P2ry12-CreER* mice were generated by CRISPR-facilitated homologous recombination. The *P2ry12* coding sequence was joined to *CreER* by deletion of the *P2ry12* stop codon and in-frame 3’ insertion of *P2A-CreER*. The P2A fusion peptide contains a 5’ GSG motif, which has been shown to significantly increase fusion protein cleavage efficiency (Szymczak-Workman et al., 2012). *P2ry12-CreER* mice were generated by pronuclear injection of the *P2ry12-CreER* targeting construct and a Cas9+gRNA expression plasmid (Biocytogen) into fertilized mouse eggs, followed by adoptive embryo transfer. Founder mice and progeny were screened by PCR, and proper genomic integration was verified by Southern blot and genomic sequencing. Recombination was induced by three doses of tamoxifen dissolved in corn oil, administered by oral gavage every other day (150uL of 20mg/mL). For embryonic mouse inductions, pregnant dams were given tamoxifen (150uL of 20mg/mL) on E13.5, E15.5 and E17.5, for a total of three gavage injections. All mouse work was performed in accordance with UCSF Institutional Animal Care and Use Committee protocols. The *P2ry12-CreER* mouse line will be deposited at Jackson Labs and at the Mutant Mouse Resource and Research Center (MMMRC).

### Histology and immunostaining

Mouse brains and other organs were harvested following transcardial perfusion with 20 mL cold PBS and 20 mL cold 4% formaldehyde. Tissue was fixed overnight at 4 degrees in 4% formaldehyde, followed by overnight incubation in 30% sucrose. Samples were embedded (Tissue Plus O.C.T. Compund, Fisher Scientific) and sectioned at 40um for adult tissues, and 20um for embryonic tissues. Sections were immunostained using a blocking/permeabilization buffer of PBS containing 2% BSA, 5% donkey serum and .5% TritonX-100. Primary and secondary antibodies were diluted in PBS containing 1% BSA and .25% TritonX-100. Secondary antibodies conjugated to Alexa fluorophores were used at 1:300 (Jackson ImmunoResearch). Primary antibodies used included: aSMA-647 (mouse monoclonal sc-32251, Santa Cruz Biotechnology, used at 1:150); Cd11b-PECy7 (BD Biosciences, used at 1:100); Cd45-APC (BD Biosciences, used at 1:100); ERTR7 (rat monoclonal abcam ab51824, used at 1:300); EGFP (chicken polyclonal, abcam ab13970 used at 1:300); TdTomato (rat monoclonal, Chromotek, 5F8, used at 1:500); TdTomato (rabbit polyclonal, Living Colors DsRed Polyclonal #632496, used at 1:1000); P2ry12 (rabbit polyclonal, kindly provided by Dr. David Julius, used at 1:1000); Iba1 (Goat, Novus NB100-1028, used at 1:300); Olig2 (Rabbit polyclonal, Millipore, used at 1:300); NeuN (mouse, Millipore MAB377, used at 1:100); Sox9 (goat, R&D System AF3075, used at 1:300); Lyve1 (Rabbit polyclonal, Abcam ab14917, used at 1:300); HA (Rabbit monoclonal, Cell Signaling C29F4, used at 1:300). Cell counting in Figure 1D was performed on three mice, using three separate images taken from the spinal cord, cerebellum, cortex and hippocampus of each mouse.

### Flow cytometry

Microglial flow cytometry was performed using a percol gradient isolation strategy. Single-cell suspensions were prepared and centrifuged over a 30%/70% discontinuous Percoll gradient (GE Healthcare) and mononuclear cells were isolated from the interface. Flow cytometry was performed on a FACS Aria III using the FACSDiva 8.0 software (BD Biosciences) and was analyzed using FlowJo v10.6.1. TdTomato+ cells were analyzed for CD45 and CD11b expression following initial gatings for forward scatter, singlets and live cells. Blood smears and flow cytometry were performed on adult mice following three tamoxifen inductions as described above. Blood was collected from the facial vein and smears were imaged for EGFP and TdTomato expression. Prior to immunostaining, blood samples used for flow cytometry were depleted of erythrocytes using an ammonium chloride lysis buffer (.15M NH4Cl, 10mM NaHCO3, 1.2mM EDTA) wash. Blood flow cytometry was performed on a BD FACSVerse machine and analyzed using FlowJo v10.6.1.

### Ribosomal immunoprecipitations

Whole brains of *P2ry12-CreER; Rpl22-HA* and *Cx3cr1-Cre; Rpl22-HA* mice were dounce homogenized in 5mL of homogenization buffer (1% NP-40, .1M KCL, 50 mM Tris pH 7.4, .02 mg/mL cycloheximide, .012M MgCl2, RNAse inhibitors, Heparin, .5 mM DTT). Tissue lysates were centrifuged at 10k-rcf for 10 min at 4 degrees to obtain a supernatant ribosomal fraction. An 80uL input sample was taken at this point to serve as a comparison to immunoprecipitated samples. Rabbit monoclonal anti-HA antibody (Cell Signaling C29F4, 2.5uL per mL) was added to the supernatant and incubated for 4 hours at 4 degrees. Iron-linked Protein A beads were added to the ribosomal fraction/antibody mixture (40uL per mL) and incubated for an additional 4 hours before being magnetically precipitated. Immunoprecipitate/beads were washed with a high salt buffer (.3M KCL, 1% NP-40, 50mM Tris pH 7.4, .012M MgCL2, cyclohexamide, .5 mM DTT) 3 times and then eluted with 350 uL of buffer RLT (Qiagen RNeasy kit). Input and immunoprecipitated RNA was purified on column (Qiagen RNeasy kit) and frozen as aliquots at −80 degrees.

### qPCR

cDNA from ribosomal immunoprecipitations was generated using Maxima First Strand cDNA Synthesis Kit (Thermo scientific) and qPCRs were performed using SYBR Green qPCR Mastermix (Bimake) on a BioRad CFX Connect thermocycler. Quantitative comparisons between immunoprecipitations and input samples were made using delta-delta Ct normalization, with GAPDH as a normalizing control.

### Library preparation and sequencing

Total RNA quality was assessed by spectrophotometer (NanoDrop, Thermo Fisher Scientific Inc., Waltham, MA) and with an Agilent Fragment Analyzer (Agilent Technologies, Palo Alto, CA). All RNA samples had integrity scores greater than 9, reflecting high-quality, intact RNA. RNA sequencing libraries were generated using the Nugen Universal Plus mRNA-Seq kit with multiplexing primers, according to the manufacturer’s protocol (NuGen Technologies, Inc., San Carlos, CA). Fragment size distributions were assessed using Agilent’s Fragment Analyzer DNA high-sensitivity kit. Library concentrations were measured using KAPA Library Quantification Kits (Kapa Biosystems, Inc., Woburn, MA). Equal amounts of indexed libraries were pooled and single end 50 bp sequencing were performed on the Illumina HiSeq 4000 at the UCSF Center for Advanced Technology (CAT) (http://cat.ucsf.edu).

### RNA sequencing analysis

Raw reads were pseudo-mapped to the mouse transcriptome using Salmon 0.13.1 (Patro et al., 2017). Quantified transcripts were passed to Tximport 1.12 for gene-level summarization and differential expression analysis was performed with DESeq2 1.24, using alpha = 0.05 (Love et al., 2015) (Love et al., 2014) . Genes with total read counts below 20 in all samples were eliminated from the analyses. Data has been submited to the Gene Expression Omnibus (GEO) repository for datasets, the accension number for this dataset is: GSE138333.

### Imaging

Confocal images were taken using a motorized Zeiss 780 upright laser scanning confocal microscope with a 34 detector array with a water immersion Zeiss Plan Apochromat 20X/1.0 D=0.17 Parfocal length 75 mm (Zeiss, Germany). Images for Figure 2 and Figure 3C were taken with a Zeiss Axio Imager.Z2 epifluorescent microscope.

## Acknowledgements

We would like to thank UCSF genomics core members Andrea Barczak, Matthew Aber, and Joshua Rudolph for their help in generating ribosomal immunoprecipitation libraries for our microglial transcriptional profiling experiments. We would also like to thank Erik Chow, Kaitlin Chaung and Derek Bogdanoff at the UCSF Center for Advanced Technology (http://cat.ucsf.edu) for their help with high-throughput sequencing. We would also like to thank Marie La Russa, Daniel Bayless and Joseph Knoedler for helpful manuscript editing suggestions.

